# Dynamic analysis of Q-fever transmission among cattle in the Tropical Savannah Grassland zone of Ghana

**DOI:** 10.1101/2024.11.15.623876

**Authors:** Dominic Otoo, Kennedy Mensah, Eugene Adjei, Baaba Abassawah Danquah, Hawa Adusei, Razak G. Chuaya

## Abstract

Livestock morbidity and death from Q-fever have been high, endangering local farmers’ livelihoods and affecting food security in Ghana. It is essential to understand the transmission dynamics of Q-fever to protect both the health of the animals and the main source of income for the community. A non-linear ordinary differential equation incorporating a vaccinated compartment was formulated and analyzed to gain insights into the spread of Q-fever. Routh Hurwitz criterion and Lyapunov function were used respectively to analyze the local and global stability of the disease-free equilibrium (*Q*_0_). We analyzed the behavior of the model compartments and discovered that many key factors significantly influence the persistence or eradication of Q-fever. Increased vaccination rates decrease the susceptible livestock while increasing the vaccinated livestock, potentially reducing the risk of outbreaks and limiting the spread of infections. A higher recovery rate leads to a quicker recovery, which aids in epidemic control by boosting population immunity and reducing the infectious time. The infection level rises when *R*_0_ *>* 1, indicating a typical transcritical bifurcation behavior, but this growth stays steady and does not result in unbounded advancement.

**Author summary:** Q-fever presents considerable health hazards to livestock in Ghana’s Tropical Savannah Grassland, adversely affecting local farmers’ income and food security in regions such as North Tongu municipality. To elucidate the transmission dynamics of the disease and safeguard both animal health and the community’s principal economic resource, we proposed a mathematical model employing non-linear differential equations that incorporate a vaccine compartment. This model enables the evaluation of how parameters such as vaccination and recovery rates influence the transmission of Q-fever in livestock. Our findings indicate that increased vaccination rates may diminish the population of susceptible livestock, hence reducing the possibility of outbreaks. In addition, a rapid recovery rate not only diminishes the duration of infectiousness in livestock but also enhances herd immunity, assisting in the containment of possible epidemics. The proposed model indicates that as *R*_0_ exceeds 1, the infection level rises, displaying transcritical bifurcation behavior. However, this rise stabilizes and prevents uncontrolled spread. These findings emphasize the value of vaccination and recovery techniques in controlling and possibly eliminating Q-fever in cattle, which would ultimately help Ghanaian farming communities remain sustainable.

## 1 Introduction

Q-fever is a zoonosis disease with active infection cases registered in most countries. Coxiella (C.) burnetii is the causative agent with a persistent infection risk in livestock. The infection is typically asymptomatic, often remaining misdiagnosed in livestock until adverse pregnancy results manifest within a herd [20]. Q-fever is found worldwide, with a higher prevalence in areas where livestock farming is common [10]. It has several different hosts including livestock, companion and wild animals [21, 22] and is only an intracellular bacterium.

The World Health Organisation (WHO) states that Rift Valley fever, Q-fever, and brucellosis are neglected zoonotic illnesses that are vulnerable to underreporting and incorrect diagnosis [12, 13]. Clinical symptoms and laboratory tests are the basis for Q-fever diagnosis, and treatment involves antibiotic therapy. Prevention measures include pasteurization of dairy products, proper disposal of animal waste, and the use of personal protective equipment when working with animals [9, 15, 16].

The mathematical modeling of infectious diseases, along with its analytical and numerical analysis, has significantly enhanced the comprehension of disease dynamics [2–4, 8]. Deterministic approaches have proven highly effective in developing realistic epidemiological models and are increasingly gaining recognition [8]. [6] pointed out that epidemic models are frequently used to clarify the mechanism of disease transmission, identify the best control measures to use, and calculate the most efficient amount of money to spend on fighting infectious diseases. To explore the evolution of the Influenza epidemic in Morocco, an SEIS compartmentalized model was formulated taken into account the seasonality of the parameter (infection, recovery and intervention rates) by [7]. Analysis of sensitivity and uncertainty was conducted to ascertain the most influential parameter and relationship between the different parameters of the model respectively. [14] examined a deterministic model for the co-infection of HPV with cervical cancer and HIV with AIDS illnesses. Using the basic reproduction number as a basis, they were able to establish the endemic and disease-free equilibrium’s local and global stability. The important parameters in their model were subjected to sensitivity analysis to determine their relative importance and possible influence on the dynamics of HIV and HPV transmission independently. A deterministic compartmental model was constructed by [4] to investigate the dynamics of infectious disease propagation while accounting for a vigilant human compartment. To ascertain whether the disease-free equilibrium was stable and whether an endemic equilibrium existed, they used a Lyapunov function.

The study of [2] presented a deterministic model to explain the potential route of Q-fever illness transmission in livestock. They employed a matrix-theoretic technique and a Lyapunov function to determine the model’s local and global asymptotic behaviour. Their sensitivity analysis shows during an outbreak, removing sick cattle from farms or culling them will lessen the influence of the animals’ exposure rate, which will lower the amount of secondary illnesses and bacterial load in the surrounding air.

A mathematical model for the spread of Q-fever in cattle herds in the United Kingdom was formulated by [23]. Regarding farm demographics and environmental impacts, their model presented the disease’s within-herd infection cycle. The results of [23] show that the outcomes of the suggested model are consistent with those of in silico investigations. It was established that the model they formulated may be used as mathematical evidence to evaluate different approaches to managing the dynamics of a Q-fever infection.

Although Q-fever has been documented in numerous regions globally, data regarding the disease in Ghana is limited.

In 2020, livestock in Ghana contributed approximately 4.4 billion Ghana cedis, equivalent to around 720.6 million USD, to the nation’s gross domestic product (GDP) [18]. The primary agricultural activity in the Tropical Savannah Grassland zone of Ghana (North Tongu municipality), is cattle rearing. However, due to little attention and negligence, Q-fever is wreaking havoc on animal lives in this community. Consequently, it is essential to understand the propagation dynamics of the disease and develop a mathematical model of Q-fever to mitigate its dissemination.

In our model, the population is subdivided into five compartments, where a confirmation of the biological realism of the model is attained. The remaining study is organized as: Section 2 presents the model’s derivation and description, Section 3 talks about the; the Basic reproduction number, disease-free equilibrium, and its stability analysis. Section 4 presents the numerical results, and the last section presents the bifurcation analysis. Finally, we conclude and assess the insights this work adds to the literature on Q-fever propagation dynamics.

## 2 Model formulation and description

Figure 1 represents the transfer diagram of individual compartment of Q-fever in the Tropical Savannah Grassland zone of Ghana. The model is divided into susceptible (*S*_*A*_), vaccinated (*V*_*A*_), exposed (*E*_*A*_), infected (*I*_*A*_), and recovered (*R*_*A*_) populations. It is assumed that the susceptible individuals are recruited into the population at a per capita rate of Λ and the class increases as a result of birth and immigration. The sum of all compartments *N* (*t*), is denoted as;

**Fig 1.**
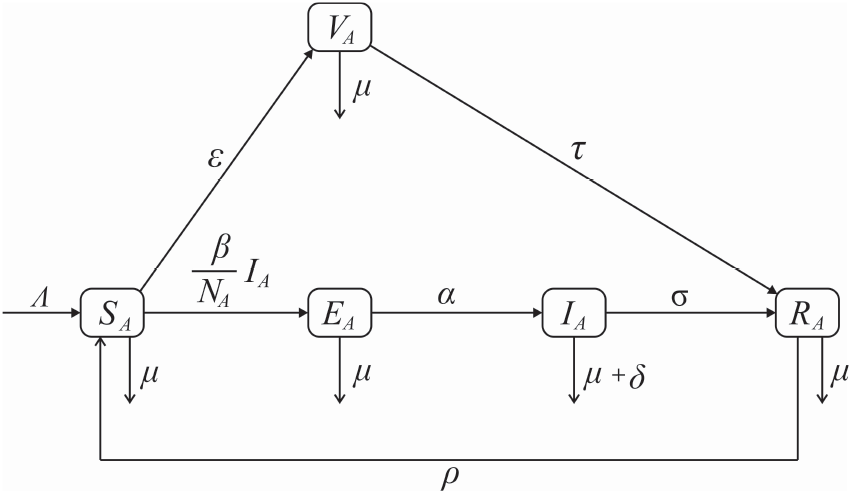
Transfer diagram of Q-fever propagation dynamics in a cattle population

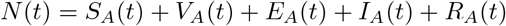

The following model equations were developed from the transfer diagram in figure 1

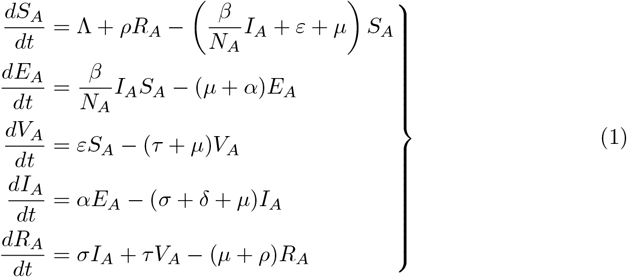

Table 1 shows the Q-fever model parameters and their descriptions as used in the model construction.

**Table 1.** Q-fever model parameters and their descriptions.

| Parameter | Description |
| --- | --- |
| $\Lambda$ | Rate of recruitment of susceptible vector |
| $\beta$ | Proportion of susceptible vector population that are exposed |
| $\tau$ | Rate at which Vaccinated vectors recover |
| $\alpha$ | Rate at which exposed individuals get infected |
| $\sigma$ | Rate Infected Vectors Recover |
| $\delta$ | Proportion of death due to Q-fever |
| $\rho$ | Rate at which recover vector becomes susceptible |
| $\mu$ | Natural death rate |
| $\varepsilon$ | Rate Susceptible Vector get Vaccinated |

### Theorem 1

*Positivity and Boundedness*

Given a positive initial values of the system *S*_*A*_(0) *≥* 0, *E*_*A*_(0) *≥* 0, *V*_*A*_(0) *≥* 0, *I*_*A*_(0) *≥* 0, *R*_*A*_(0) *≥* 0, then the solution set of the outcome of

*S*_*A*_(*t*), *E*_*A*_(*t*), *V*_*A*_(*t*), *I*_*A*_(*t*), *R*_*A*_(*t*) of the system (1) is non-negative and bounded *∀t >* 0 wherever they exist.

**Proof 1** *From the system in (1)*,

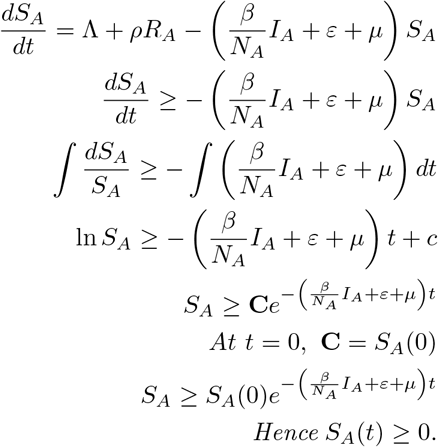

*Similar approach on the rest of the compartments gives*

*E*_*A*_(*t*) *≥* 0, *V*_*A*_(*t*) *≥* 0, *I*_*A*_(*t*) *≥* 0, *R*_*A*_(*t*) *≥* 0.

*Hence, it is clear the vector population state variables are positive and bounded*.

### Theorem 2

*Invariant Region*

The set of solution of 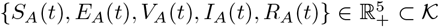.

**Proof 2** *Given the population*,

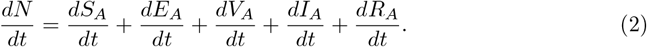

*Basic substitution of system (1) into equation 2 gives;*

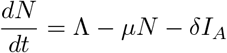

*Applying separation of variables*

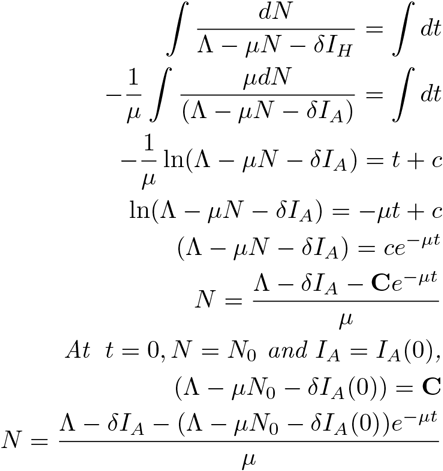

As 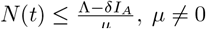.

*Hence* 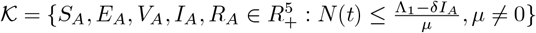 *is positively invariant for all non-negative values of t*.

## 3 Model analysis

### 3.1 Disease free equilibrium (*Q*_0_)

The disease-free equilibrium happens in the Q-fever system (1) when the infection compartments (*E*_*A*_ = *I*_*A*_ = 0). Hence

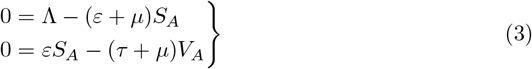

From Eq. (3), the following deductions can be made;

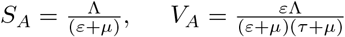

Hence,

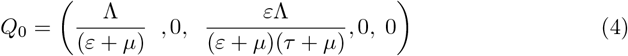

### 3.2 Basic Reproduction Number (*R*_0_)

The infection compartments of model (1) are given by;

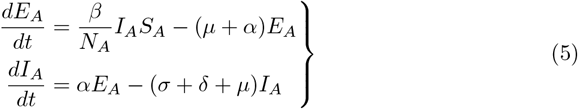

Following [2] and employing the next generation matrix on system of equations (5), matrix *F* and *V* are deduced as follows;

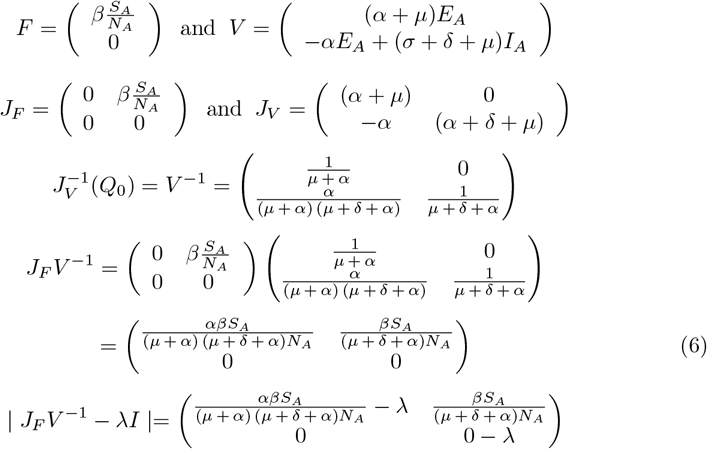

Hence, the greatest eigenvalues of *J*_*F*_ *V* ^*−*1^ gives

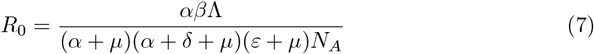

#### Theorem 3

*Local stability of Q*_0_

The point *Q*_0_ exhibits local asymptotic stability whenever *R*_0_ *<* 1 and unstable if *R*_0_ *>* 1 given system (1).

**Proof 3** *Let Q*_0_ *be the Q-fever free equilibrium point, then the linearization of system of equations (1) gives;*

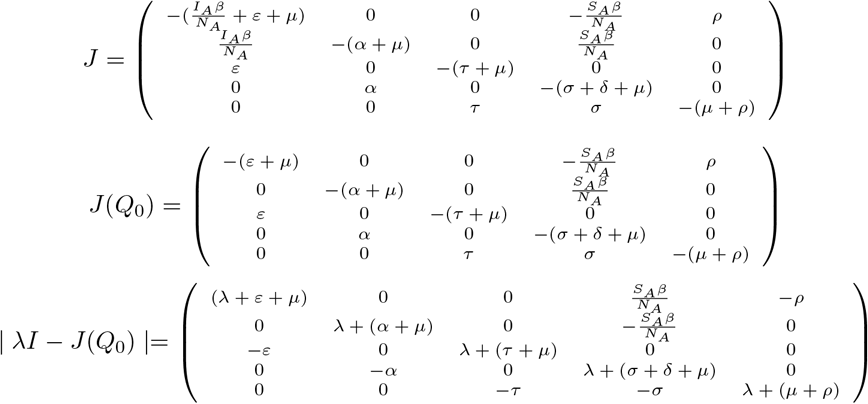

*Now* | *λI − J* (*Q*_0_) |= 0 *implies*,

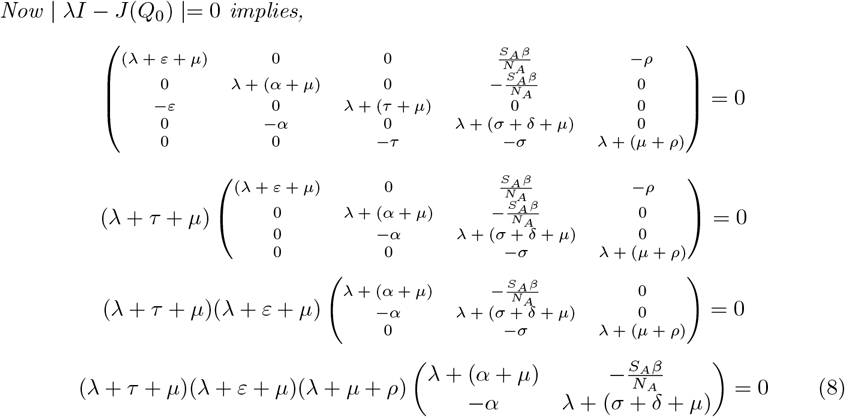

*Clearly all the λ*^*′*^*s of* (*λ* + *τ* + *μ*)(*λ* + *ε* + *μ*)(*λ* + *μ* + *ρ*) *are negative. The remaining eigenvalues are obtained from* 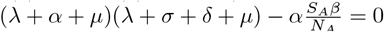.

*Let* (*α* + *μ*) = *a and* (*σ* + *δ* + *μ*) = *b*.

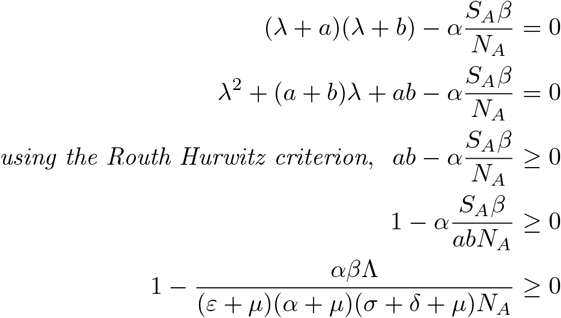

*Thus* 1 *− R*_0_ *≥*_0_, *and Q*_0_ *is local asymptotically stable*.

#### Theorem 4

*Global stability of Q*_0_

The point *Q*_0_ of system (1) is globally asymptotically stable for *R*_0_ *<* 1 and unstable for *R*_0_ *>* 1.

**Proof 4** *We employ the LaSalle Invariance Principle [25], to prove the globally asymptotic stability of Q*_0_. *Let*

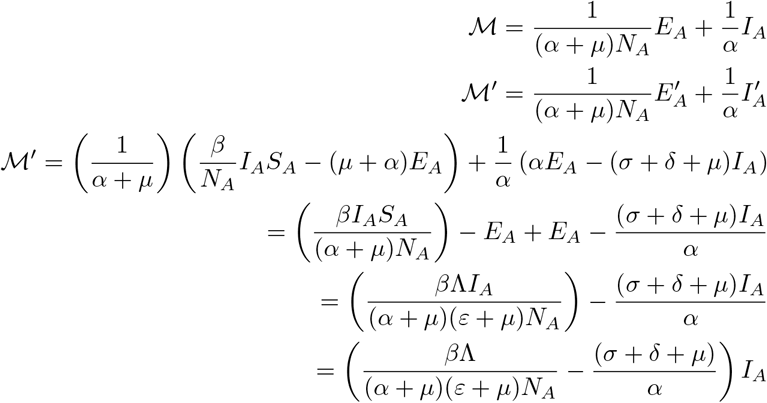

*But*

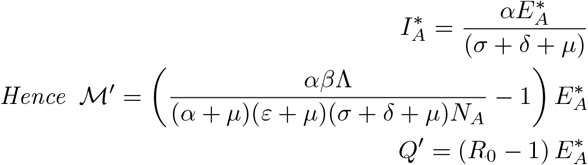

*For R*_0_ *<* 1 *and* 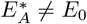. *Hence, the Lyapunov function ℳ at Q*_0_ *is globally asymptotically stable for R*_0_ *<* 1 *while unstable for R*_0_ *>* 1.

### 3.3 Existence of Q-fever endemic equilibrium point (*Q*_*p*_)

Setting the model (1) to zero and solving for the state equations, we have

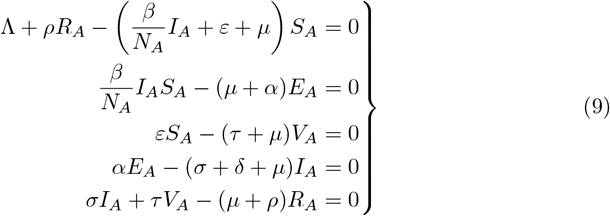

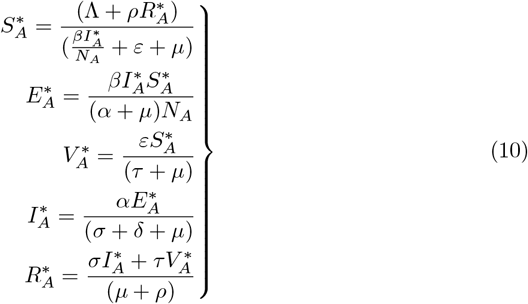

#### Theorem 5

*Local stability of Q*_*p*_

The equiibrium point *Q*_*p*_ of system (1) is locally asymptotically unstable if *R*_0_ *>* 1 and stable otherwise.

**Proof 5** *Linearizing the system (1) gives;*

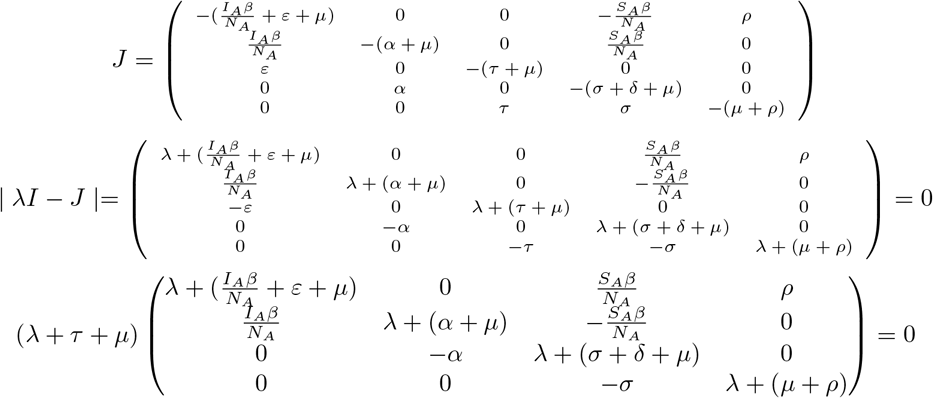

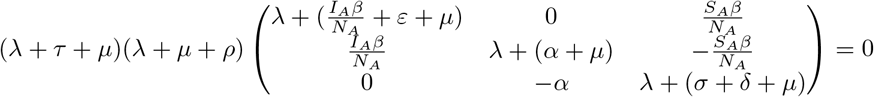

*It is trivial that the eigenvalues of* (*λ* + *τ* + *μ*)(*λ* + *μ* + *ρ*) *are negatives. The remaining eigenvalues are from*

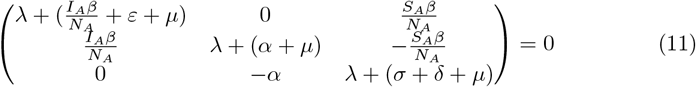

*Let* 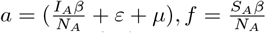, *b = (α + μ), c = (σ + δ + μ)*

*Then Eq. (11) becomes*

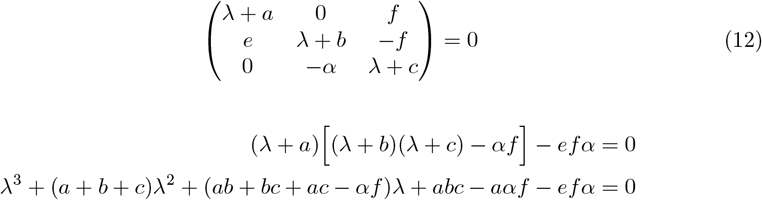

*Let p*_0_ = *abc − aαf − efα, p*_1_ = (*ab* + *bc* + *ac − αf*), *p*_2_ = (*a* + *b* + *c*).

*Routh-Hurwitz array:*

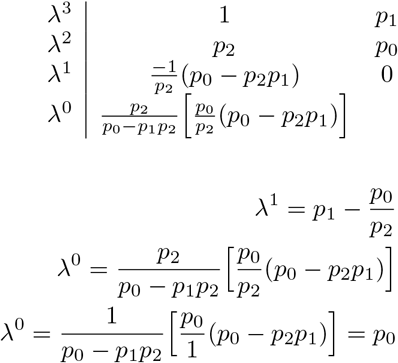

*From the array the remaining eigenvalues have negative real parts if p*_0_ *>* 0 *and p*_1_*p*_2_ *> p*_0_. *Computation in both R and Maxima software shows p*_0_ = 0.038 *and p*_1_*p*_2_ = 0.534.

*Hence Q*_*p*_ *is locally asymptotically unstable whenever R*_0_ *>* 1.

## 4 Numerical analysis and discussion

To examine key model parameters and evaluate each compartment’s dynamism, we conducted numerical simulations using parameter values of Table 2. These are simulated values and others are taken from published article [27]. After examining the model compartments’ behavior, we found that several significant variables have a considerable effect on whether Q-fever persists or is eradicated. The characteristics stated above can also provide insight into how to lower transmissions. Figure (2) provides useful epidemiological meanings by illustrating the dynamic properties of each compartment, especially when looking at equilibrium states and stability. This image illustrates how an outbreak develops through various phases and how the population in each compartment varies over time.

**Table 2.**
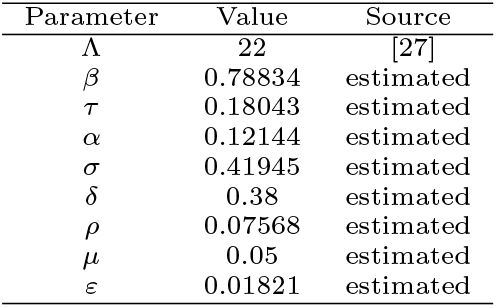
Parameters and their respective values.

The figure (3) shows how varying parameters like *β, τ, ε* can regulate infection patterns, providing insights for effective disease management approaches in livestock herds. High contact rate causes an increase in the number of both exposed and infected cattle, which spreads infection more quickly. However, when susceptible livestock become exposed or infected their numbers decrease more quickly. A higher vaccination rate lowers the susceptible population while increasing the vaccinated population, potentially lowering the likelihood of outbreaks and limiting disease spread. The greater the recovery rate, the faster the recovery, building up immunity in the population and potentially decreasing the infectious time, which helps to manage the epidemic.

**Fig 2.**
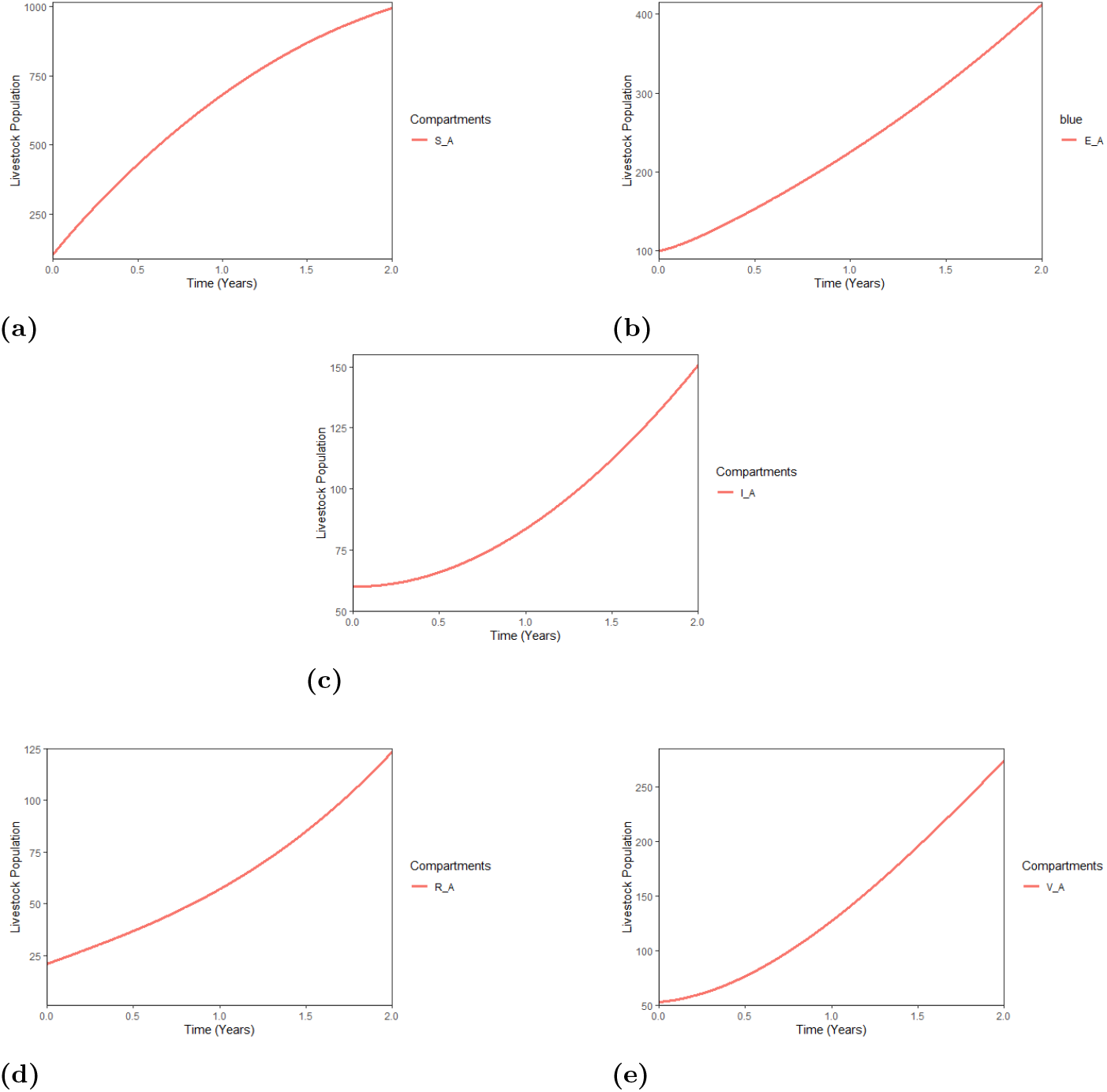
Phase plot for *SV EIR* model with time.

**Fig 3.**
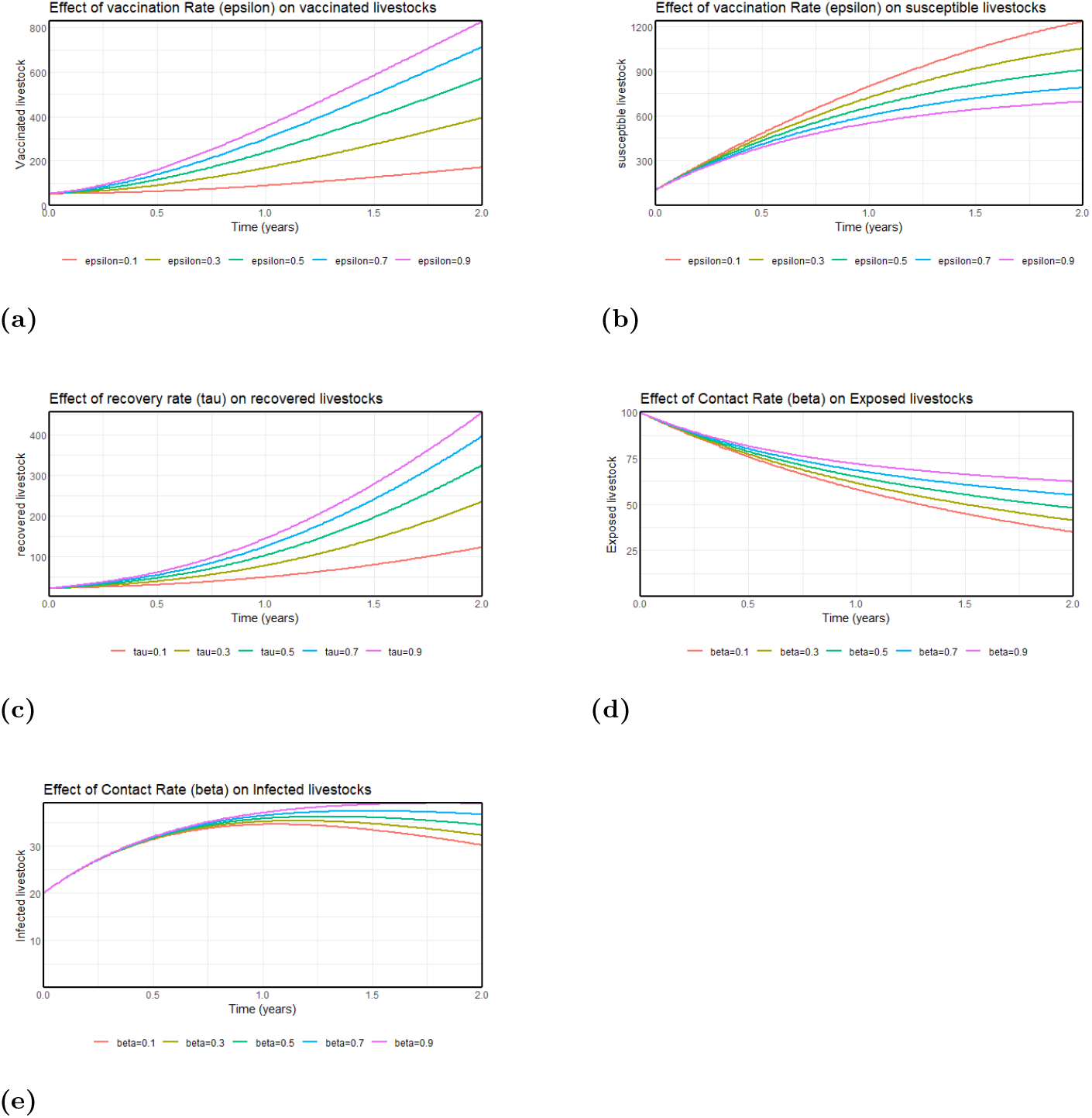
Transmission trajectories based on modifying pertinent variables.

Simulations of the rate of transmission in exposed and infected livestock at various *R*_0_ rates can be observed in Figure 4. For all values of *R*_0_, the number of exposed cattle (Fig. 4 **a**) primarily rises quickly. The number of exposed livestock begins to reduce with time, suggesting that the outbreak is peaking and beginning to decline. For all values of *R*_0_, the number of infected livestock originally rises slowly (Fig. 4 **b**).

**Fig 4.**
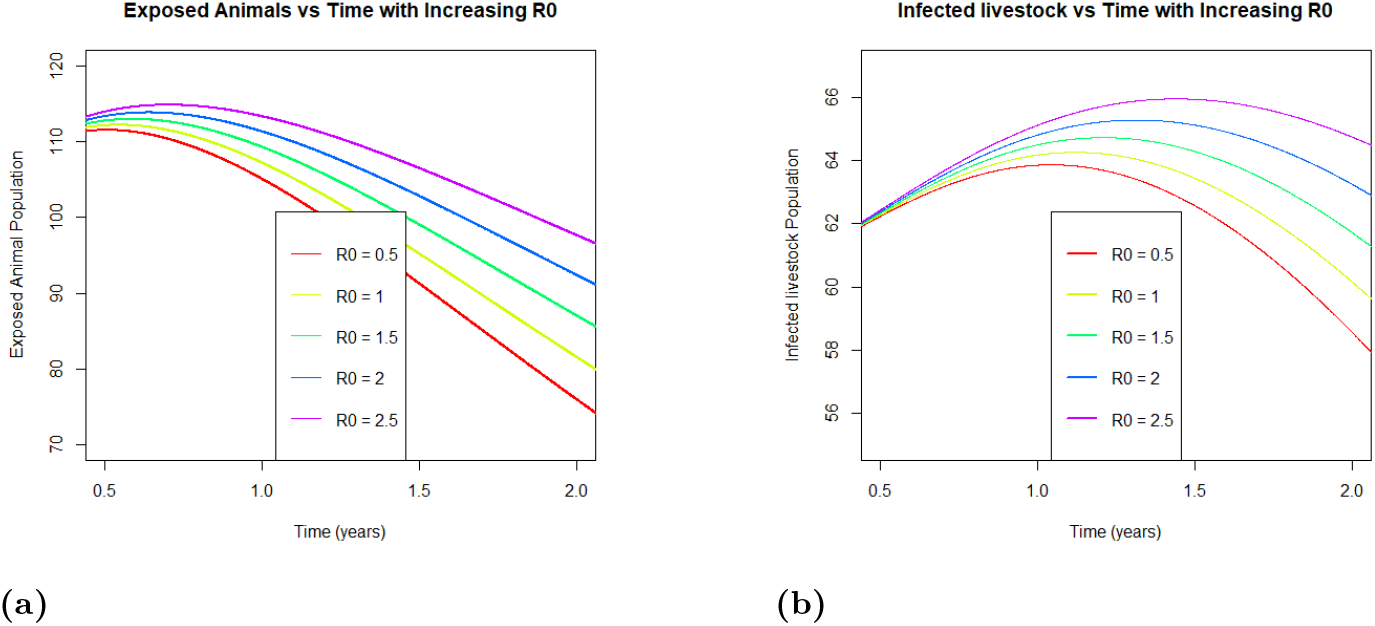
Effect of an increase in *R*_0_ on exposed and infected livestocks

The number of infected cattle begins to rise more rapidly than the number of exposed animals, with the peak of infected cattle occurring later than that of the exposed population. A greater value of *R*_0_ is associated with a higher peak outbreak and more rapid initial emergence of exposed and infected livestock. This emphasizes the significance of regulating *R*_0_ by vaccination to prevent or lessen the impact of outbreaks.

## 5 Bifurcation Analysis

The central manifold analytical method of Castillo-chavez and Song [17] will be considered to study the kind of bifurcation the Q-fever model exhibits. Consider the system of differential equation;

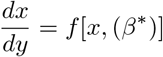

where the bifurcation parameter is *β* and *f* represents a differentiable continuous function which appears at least two times in *x*. The coefficients *a* and *b* can be calculated by making use of the fact that;

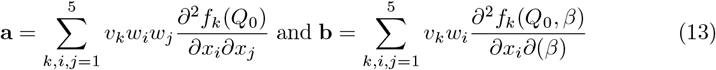

This, when combined with the conditions of [17], show the existence of a bifurcation in the Q-fever model. Now, considering the left eigenvectors of system (1),

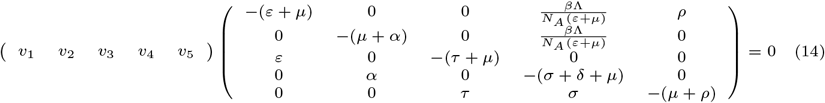

The following equations result from (14):

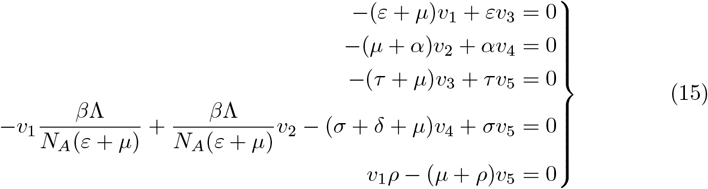

Applying the right eigenvector,

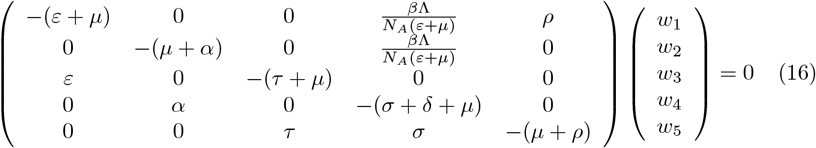

From 16, the following equations can be deducted;

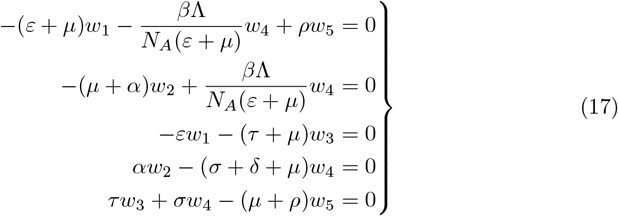

To compute the value of **a**, we use the relation (13).

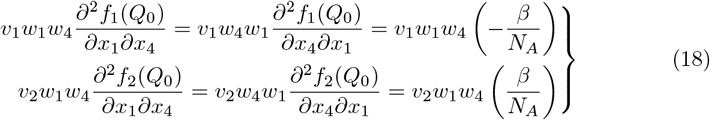

Hence

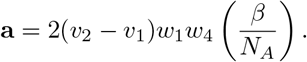

By substitution,

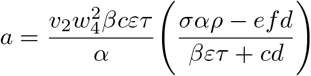

where *c* = (*τ* + *μ*), *d* = (*μ* + *ρ*), *e* = (*σ* + *δ* + *μ*), *f* = (*μ* + *α*).

Since *efd > σαρ, v*_2_ *>* 0, showing that *a <* 0.

The computation of **b** is given by;

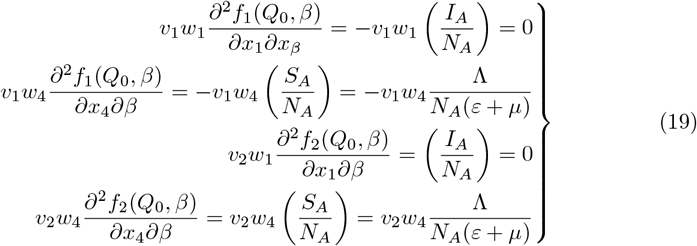

Hence,

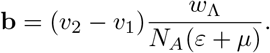

Simplify, we have

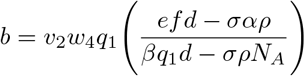

where 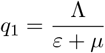 and *c, d, e, f* are as stated above.

But *efd > σαρ, σρN*_*A*_ *> βq*_1_*d, v*_2_ *>* 0, *w*_4_ *<* 0 and hence *b >* 0.

From the endemic equilibrium, 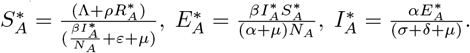

By direct substitution,

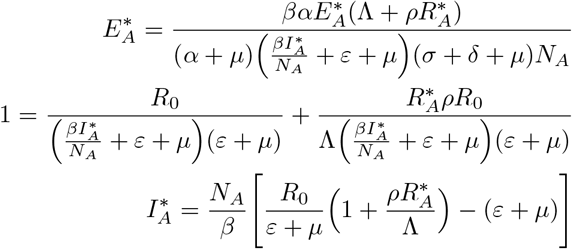

*Q*_0_ is usually stable for *R*_0_ *<* 1 when the condition *a <* 0 is met. This means that any disturbances around zero levels of infection will decay, eventually bringing the system back to *Q*_0_. For *R*_0_ *>* 1, the system appears to have a stable positive *Q*_*p*_, according to the condition *b >* 0. The infection level increases when *R*_0_ *>* 1, in line with a typical transcritical bifurcation behavior [25], but this growth remains constant and does not lead to unlimited advancement. A clear demonstration of the phase transition among disease-free and endemic equilibrium, which is controlled by a basic reproduction number, *R*_0_, is shown in Figure 5. It highlights the significance of maintaining *R*_0_ below 1 to prevent the disease from spreading.

**Fig 5.**
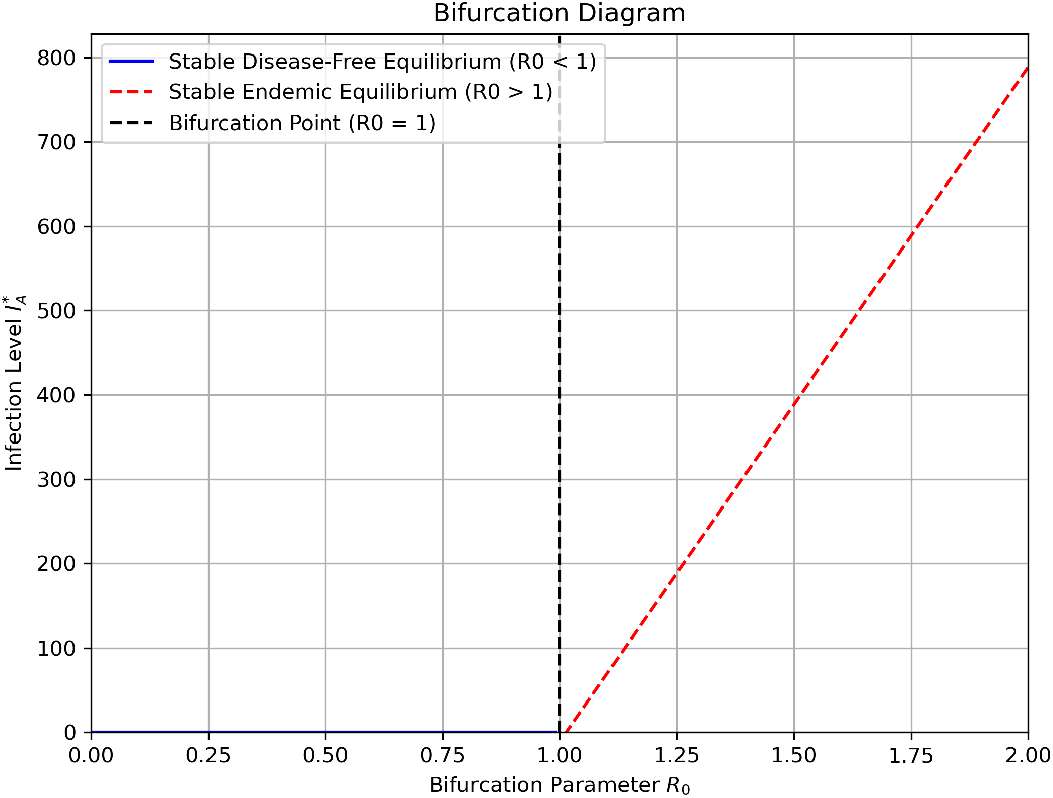
Forward (transcritical) bifurcation

## 6 Conclusions

In this study, a deterministic mathematical model was formulated to describe the transmission dynamics of Q-fever in the Tropical Savannah Grassland zone of Ghana. The formulated model, using a system of non-linear ordinary differential equations, was analyzed to understand the propagation dynamics of the disease and to mitigate its dissemination. The model satisfies fundamental biological relevance and reliability, including positivity, boundedness, and the existence of an invariant region. Routh Hurwitz criterion and Lyapunov function were used respectively to analyze the local and global asymptotic stability of the disease-free equilibrium (*Q*_0_). We analyzed the behavior of the model compartments and discovered that many critical factors significantly influence the persistence or eradication of Q-fever. It can be observed that the higher the value of *R*_0_, the faster the initial increase in exposed and infected livestock and the higher the peak of the outbreak. The infection level rises when *R*_0_ *>* 1, indicating a typical transcritical bifurcation behavior, but this growth stays steady and does not result in unbounded advancement. Further analysis incorporating environmental or seasonal factors and optimal control strategies could enhance the model’s applicability to real-world scenarios, supporting more effective, data-driven decision-making in livestock health management.

## Data Availability Statement

The parameter values (data) used to support the findings of this study have been described in Section 2 and stated in Section 4. Some of the parameter values are simulated values and others are taken from published article and are cited at relevant places within the text as reference [27].

## Acknowledgments

We appreciate the editor’s and reviewers’ insightful feedback. Also, the authors are grateful to the Veterinary Services Directorate, North Tongu, for making the data available for the purpose of this study.

## CRediT authorship contribution statement

Dominic Otoo: Conceptualization, Methodology, Formal analysis, Supervision. Kennedy Mensah: Formal analysis, Validation, Writing – original draft, Writing – review & editing. Eugene Adjei: Methodology, Formal analysis. Baaba Abassawah Danquah: Methodology, Conceptualization. Hawa Adusei: Methodology, Formal analysis. Charles Sebil: Software, Data curation, Conceptualization.Razak G. Chuaya: Conceptualization, Methodology, Software.

## References

1. Courcoul, Aurélie and Hogerwerf, Lenny and Klinkenberg, Don and Nielen, Mirjam and Vergu, Elisabeta and Beaudeau, François (2011). Modelling effectiveness of herd level vaccination against Q fever in dairy cattle. Veterinary research, 42(1), 1–9.

2. Asamoah, Joshua Kiddy K and Jin, Zhen and Sun, Gui-Quan and Li, Michael Y and others (2020). A deterministic model for Q fever transmission dynamics within dairy cattle herds: using sensitivity analysis and optimal controls. Computational and Mathematical methods in Medicine, 2020.

3. Asamoah, Joshua Kiddy K and Jin, Zhen and Sun, Gui-Quan (2021). Non-seasonal and seasonal relapse model for Q fever disease with comprehensive cost-effectiveness analysis. Results in Physics, 22, 103889.

4. Obabiyi, Olawale and Olaniyi, Samson (2019). Global stability analysis of malaria transmission dynamics with vigilant compartment.

5. Wielders, Cornelia C. H. and van Loenhout, Joris A. F. and Morroy, Gabriëlla and Rietveld, Ariene and Notermans, Daan W. and Wever, Peter C. and Renders, Nicole H. M. and Leenders, Alexander C. A. P. and van der Hoek, Wim and Schneeberger, Peter M. (2015). Long-Term Serological Follow-Up of Acute Q-Fever Patients after a Large Epidemic. PLOS ONE, 10(), 1–15.

6. Otoo, Dominic and Abeasi, Isaac Odoi and Osman, Shaibu and Donkoh, Elvis Kobina (2021). Mathematical modeling and analysis of the dynamics of hepatitis b with optimal control. Commun. Math. Biol. Neurosci., 2021, Article–ID.

7. Sara, Bidah and Omar, Zakary and Abdessamad, Tridane and Mostafa, Rachik and Hanane, Ferjouchia (20202). Parameters’ estimation, sensitivity analysis and model uncertainty for an influenza a mathematical model: case of morocco. Commun. Math. Biol. Neurosci., 2020, Article–ID.

8. Baba, I. A., Hincal, E., & Rihane, F. A. (2024). Exploring the dynamics of monkeypox: a fractional order epidemic model approach. Journal of Applied Mathematics and Computational Mechanics, 23(1), 32–44.

9. Cho YS, Park JH, Kim JW, et al. (2023). Current Status of Q Fever and the Challenge of Outbreak Preparedness in Korea: One Health Approach to Zoonoses. J Korean Med Sci., 38(24), e197.

10. Deressa, F. B., Kal, D. O., Gelalcha, B. D., & Magalhães, R. J. S. (2020). Seroprevalence of and risk factors for Q fever in dairy and slaughterhouse cattle of Jimma town, South Western Ethiopia. BMC Veterinary Research, 16, 1–10.

11. Aljafar, A., Salem, M., Housawi, F., Zaghawa, A., & Hegazy, Y. (2020). Seroprevalence and risk factors of Q-fever (C. burnetii infection) among ruminants reared in the eastern region of the Kingdom of Saudi Arabia. Tropical Animal Health and Production,52, 2631–2638.

12. Schwarz, N. G., Loderstaedt, U., Hahn, A., Hinz, R., Zautner, A. E., Eibach, D., & Frickmann, H. (2017). Microbiological laboratory diagnostics of neglected zoonotic diseases (NZDs). Acta tropica., 165, 40–65.

13. Kanouté, Y.B.; Gragnon, B.G.; Schindler, C.; Bonfoh, B.; Schelling, E. (2017). Epidemiology of brucellosis, Q Fever and Rift Valley Fever at the human and livestock interface in northern Côte d’Ivoire. Acta Trop., 165, 66–75.

14. Gurmu, ED and Bole, BK and Koya, PR (2020). Mathematical model for co-infection of HPV with cervical cancer and HIV with AIDS diseases. International journal of scientific research in mathematical and statistical sciences, 7(2).

15. Salman, M., & Steneroden, K. (2022). Important Zoonotic Diseases of Cattle and Their Prevention Measures. In Zoonoses: Infections Affecting Humans and Animals, pp. 1–22. Cham: Springer International Publishing.

16. Bond, K. A., Vincent, G., Wilks, C. R., Franklin, L., Sutton, B., Stenos, J., … & Firestone, S. M. (2016). One Health approach to controlling a Q fever outbreak on an Australian goat farm. Epidemiology & Infection,, 144(6), 1129–1141.

17. Bakare, E. A., & Nwozo, C. R. (2017). Bifurcation and sensitivity analysis of malaria–schistosomiasis co-infection model. International Journal of Applied and Computational Mathematics, 3, 971–1000.

18. Doris D. S. (2024). Contribution of livestock to Gross Domestic Product (GDP) in Ghana from 2013 to 2022. https://www.statista.com/statistics/1272321/annual-contributions-of-livestock-to-gdp-in-ghana.

19. Asamoah, Joshua Kiddy K and Oduro, Francis T and Bonyah, Ebenezer and Seidu, Baba. (2017). Modelling of rabies transmission dynamics using optimal control analysis. Journal of Applied Mathematics, 1(2017), 2451237.

20. Ullah, Qudrat and Jamil, Tariq and Saqib, Muhammad and Iqbal, Mudassar and Neubauer, Heinrich. (2022). Q Fever—A Neglected Zoonosis. Microorganisms, 10(8). DOI = 10.3390/microorganisms10081530

21. Abdel-Moein K. A., & Hamza D. A. (2017). The burden of Coxiella burnetii among aborted dairy animals in Egypt and its public health implications.Acta Trop., 166, 92–95. doi:10.1016/j.actatropica.2016.11.011

22. Eldin, C., Mélenotte, C., Mediannikov, O., Ghigo, E., Million, M., Edouard, S., Mege, J.L., Maurin, M. and Raoult, D. (2017). From Q fever to Coxiella burnetii infection: a paradigm change. Clinical microbiology reviews, 30(1), 115–190.

23. Patsatzis, D. G., Wheelhouse, N., & Tingas, E. A. (2022). Modelling the Transmission of Coxiella burnetii within a UK Dairy Herd: Investigating the Interconnected Relationship between the Parturition Cycle and Environment Contamination Veterinary Sciences, 9(10), 522.

24. Li, J., Blakeley, D., & Smith, R. J. (2011). The failure of R0. Computational and mathematical methods in medicine, 2011, 527610. 10.1155/2011/527610

25. Shuai, Zhisheng and van den Driessche, P. (2013). Global Stability of Infectious Disease Models Using Lyapunov Functions. SIAM Journal on Applied Mathematics, 4(73), 1513–1532. 10.1137/120876642

26. Han, X., Liu, H., Lin, X., Wei, Y., & Ming, M. (2022). Dynamic analysis of a VSEIR model with vaccination efficacy and immune decline. Advances in Mathematical Physics, 2022(1), 7596164.

27. Asamoah, J. K. K., Okyere, E., Yankson, E., Opoku, A. A., Adom-Konadu, A., Acheampong, E., & Arthur, Y. D. (2022). Non-fractional and fractional mathematical analysis and simulations for Q fever. Chaos, Solitons & Fractals, 156, 111821.

